# Genetic Subtypes of Smoldering Multiple Myeloma are associated with Distinct Pathogenic Phenotypes and Clinical Outcomes

**DOI:** 10.1101/2021.12.10.471975

**Authors:** Mark Bustoros, Shankara Anand, Romanos Sklavenitis-Pistofidis, Robert Redd, Eileen M. Boyle, Benny Zhitomirsky, Andrew J Dunford, Yu-Tzu Tai, Selina J Chavda, Cody Boehner, Carl Jannes Neuse, Mahshid Rahmat, Ankit Dutta, Tineke Casneuf, Raluca Verona, Efstathis Kastritis, Lorenzo Trippa, Chip Stewart, Brian A. Walker, Faith E. Davies, Meletios-Athanasios Dimopoulos, Leif Bergsagel, Kwee Yong, Gareth J. Morgan, François Aguet, Gad Getz, Irene M. Ghobrial

## Abstract

Smoldering multiple myeloma (SMM) is a precursor condition of multiple myeloma (MM) with significant heterogeneity in disease progression. Existing clinical models of progression risk do not fully capture this heterogeneity. Here we integrated 42 genetic alterations from 214 SMM patients using unsupervised binary matrix factorization (BMF) clustering and identified six distinct genetic subtypes. These subtypes were differentially associated with established MM-related RNA signatures, oncogenic and immune transcriptional profiles, and evolving clinical biomarkers. Three subtypes were associated with increased risk of progression to active MM in both the primary and validation cohorts, indicating they can be used to better predict high and low-risk patients within the currently used clinical risk stratification model.

---

Multiple Myeloma (MM) is an incurable plasma cell malignancy with significant inter- and intra-patient heterogeneity. It is almost always preceded by the asymptomatic precursor stages monoclonal gammopathy of undetermined significance (MGUS) and smoldering multiple myeloma (SMM). Approximately 1.5% of MGUS patients will progress to MM per year, while SMM patients have a higher overall progression risk of 10% per year^1,2^. Like MM, SMM is a heterogeneous condition—some patients have over a 50% risk of progression within two years, while others have more MGUS-like disease that progresses slowly^3^.

Several risk stratification models exist to help clinicians differentiate patients with high risk of progression to active myeloma from those for whom a ‘watchful waiting’ approach is appropriate. The existing models rely solely on clinical measurements, many of which are indicators of tumor burden and universal biomarkers of MM for risk stratification. These models, however, do not fully partition progressors from non-progressors, and patients classified as low- or intermediate-risk still progress to active MM and have a 2-year progression risk of 6% and 18%, respectively (compared to 44% for high-risk patients)^3^, which warrants a more accurate models that also represent the molecular heterogeneity in MM. We recently showed that genomic alterations in mitogen-activated protein kinase (MAPK) and DNA repair pathways or *MYC* are independently predictive of progression risk^4^. While these genomic biomarkers improved upon the clinical models, they represent only a few alterations that do not capture the full extent of genetic heterogeneity in SMM.

Multiple myeloma is characterized by multiple chromosomal gains or losses, structural variations, driver single nucleotide variations (SNVs)^5-7^, and other structural alterations involving known oncogenes^8^. The IgH translocations and copy number alterations (CNAs) are considered early events in the pathogenesis of MM, while other CNAs and SNVs usually occur later during clonal evolution, providing more proliferative capacity to the tumor cells^4^. Multiple SMM studies have shown that CNAs, including whole chromosome duplications and arm-level losses or gains, are the most common events, followed by SNVs, and then translocations^4,7,9,10^. These alterations have all been detected at the SMM stage, and we previously showed that the genomic makeup of the MM tumor clone is fully acquired by the time of SMM diagnosis in most cases^4,10-12^.. Furthermore, certain genetic alterations were found to occur more frequently together^4,7^, therefore, studying these genetic alterations as groups or networks rather than individual risk factors may improve our understanding of disease molecular groups and the overall risk stratification in SMM.

In this study, we apply an unsupervised binary matrix factorization (BMF) clustering method to identify groups of genomic alterations that tend to occur together, and show that the resulting clusters represent distinct biological and clinical subtypes in SMM

## Results

### Identification of six clusters with distinct genetic features

We leveraged DNA sequencing data from a cohort of 214 patients at the time of SMM diagnosis, with matched RNA sequencing data of 89 patients from the same cohort, and baseline clinical information for the whole cohort (**Figure 1A and Supp. Table 1**). The patients in this cohort harbored a median of 7 driver events, several of which tended to co-occur^4,6,7^, suggesting that additional analyses may reveal distinct patterns (i.e., clusters) of genetic alterations. To identify these patterns, binarized DNA features (42 driver SNVs, CNVs, and translocations) were curated for each sample representing the presence or absence of each genomic alteration. Chapuy *et al* successfully subtyped diffuse B-cell lymphoma patients with consensus non-negative matrix (NMF) factorization of numeric DNA features. We instead apply consensus BMF for this multi-omics subtyping to accommodate these binarized DNA features, appropriately model summative features that span multiple subtypes (i.e. hyperdiploidy), and handle sparse matrices (**Methods**). We identified six distinct patterns of drivers that accordingly partitioned the patient to six clusters based on their most enriched pattern. Samples in four of these clusters were hyperdiploid (more than 48 chromosomes), while those in the other two were enriched for known MM IgH translocations (**Figure 1B**).

**Figure 1.**
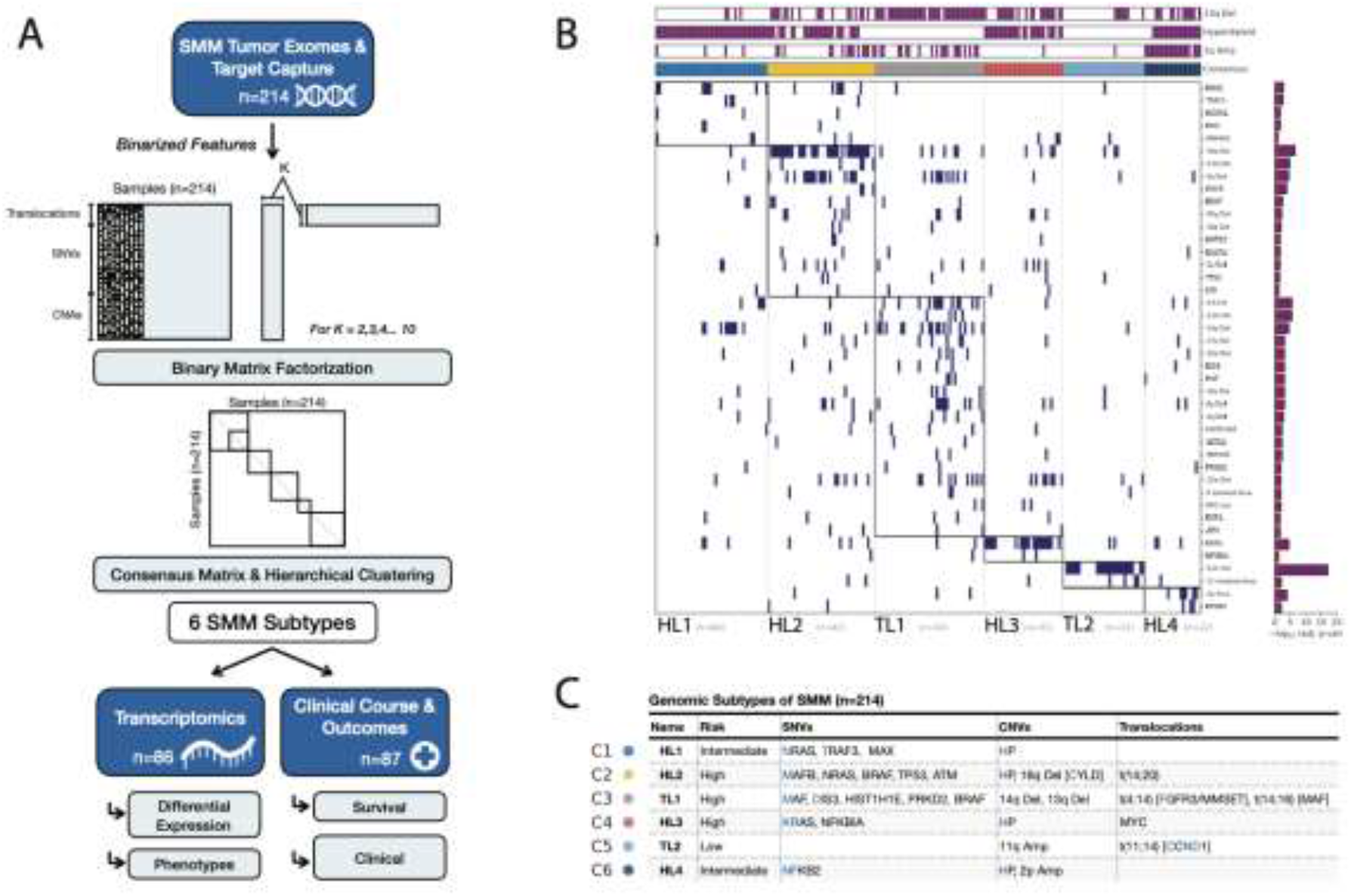
A) Flowchart of analyses performed in this study. Clusters were generated based on DNA sequencing data, then clusters were analyzed for correlations with transcriptomic and clinical data. B) The genomic profile of individual patients with smoldering multiple myeloma grouped by cluster and significant genomic features. C) Summary table including the 6 subtypes identified and enriched genetic features.

*Cluster 1:* the tumors of this cluster exhibited a hyperdiploid genotype as the primary event and were significantly enriched in *NRAS, TRAF3*, and *MAX* mutations. We named this cluster *Hyperdiploid-like1* (*HL1*). *Cluster 2:* the tumors of this cluster harbored frequent arm-level deletions, including 16q, 6q, 1p, 17p, 4q, 18q, and 20q, and the IgH translocation t(14;20), which upregulates the transcription factor *MAFB*. Moreover, mutations in *NRAS, BRAF, TP53, ATM, MAFB*, and *CDKN2C* genes were enriched in this subgroup. Hyperdiploidy was detected in 69% of the tumors in this cluster. We named this cluster *Hyperdiploid-like2* (*HL2*). The tumors of this cluster were significantly enriched in deletion(16q), which involves CYLD tumor suppressor and other genes. The presence of both hyperdiploidy and t(14;20) in the same cluster could be explained by either the small number of samples with these alterations as seen in other studies as well, or that in few cases they co-occur together. Indeed, half of patients with t(14;20) in our cohort had hyperdiploidy. *Cluster 3*: Tumors of this cluster exhibited primary events such as t(4;14), which upregulates *FGFR3* and *MMSET* genes; t(14;16), which upregulates the transcription factor *MAF*; and copy number losses of 14q, 1p, 8p, 10p, 11q, 12p, and 17p. We named this cluster *Translocation-like1* (*TL1*). This cluster was also enriched for hypodiploid tumors, defined as having fewer than 45 chromosomes (adjusted *P* = 0.04). Tumors in this cluster also harbored mutations in *BRAF, DIS3, MAF, FGFR3, PRKD2, PRDM1, IRF4*, and *HIST1H1E*. Many of these proteins and mutations in their encoding genes are essential to tumor cell survival and play roles in protein translation, secretion, and plasma cell differentiation ^7,9,13^. Differential gene expression analysis revealed that *TL1* tumors have downregulation of ribosomal proteins and the negative regulator of the MAPK pathway *TRAF2*. The upregulated genes included *WHSC1*(*MMSET*), *FGFR3, KLHL4, CCND3*, and genes involved in the endoplasmic reticulum (ER) stress response (**Figure 2A**). *Cluster 4*: this cluster comprises tumors with a hyperdiploid genotype that harbored mutations in *KRAS* and *NFKBlA* genes, and *MYC* translocations as the only significant features. We named this cluster *Hyperdiploid-like3* (*HL3*). *Cluster 5*: the tumors in this cluster mainly exhibited t(11;14), *CCND1* mutations, and gain of chromosome 11 or its long arm. We named this cluster *Translocation-like1* (*TL1*). Interestingly, this cluster had significantly lower M-protein levels and was enriched in light-chain disease compared to the other clusters (*P* < 0.001 for both). Tumors of *TL2* had 243 differentially expressed genes (q <0.1, log2FC|> 1.5; 180 upregulated, 63 downregulated), including overexpression of *CCND1, ERBB4, E2F7, E2F1, TRAK2, RBL1*, and downregulation of *DUSP4, TRAF6, PRKD3, CCDC6*, and *ZNF844*. Furthermore, this cluster had the highest expression of *CCND1* compared to the other clusters (**Figure 2B**). *Cluster 6*: this is a hyperdiploid cluster similar to *HL1*; however, its tumors are also enriched in *NFKB2* and *KLHL6* mutations and exhibit copy gains in 2p. Interestingly, copy gains of 1q were more frequent in this cluster than *HL1* and the other hyperdiploid clusters (*P* < 0.001 for both comparisons). We named this cluster *Hyperdiploid-like4* (*HL4*). Additionally, key individual genes in myeloma pathogenesis were overexpressed in tumors of specific genetic subtypes. *MCL1* was upregulated in all the genetic subgroups with the lowest expression observed in *HL1* tumors compared to the other subtypes (*P* = 0.001) (**Figure 2C**). *MYC* oncogene was also highly expressed in the four hyperdiploid clusters (*P* = 0.009, Wilcoxon Test) (**Figure 2D**). Cyclin D1 (*CCND1*) was significantly upregulated in *TL2* tumors (*P* = 0.0001), while *CCND2* was upregulated in *TL1* and *HL2* tumors compared to the rest of the genetic subtypes (*P* = 0.004) (**Figure 2E, F**). Moreover, in the four hyperdiploid clusters, we found that CCND1 expression was higher in tumors with 11q gain, while *CCND2* expression was higher in samples without 11q gain (**Supp Figure 1**).

**Figure 2.**
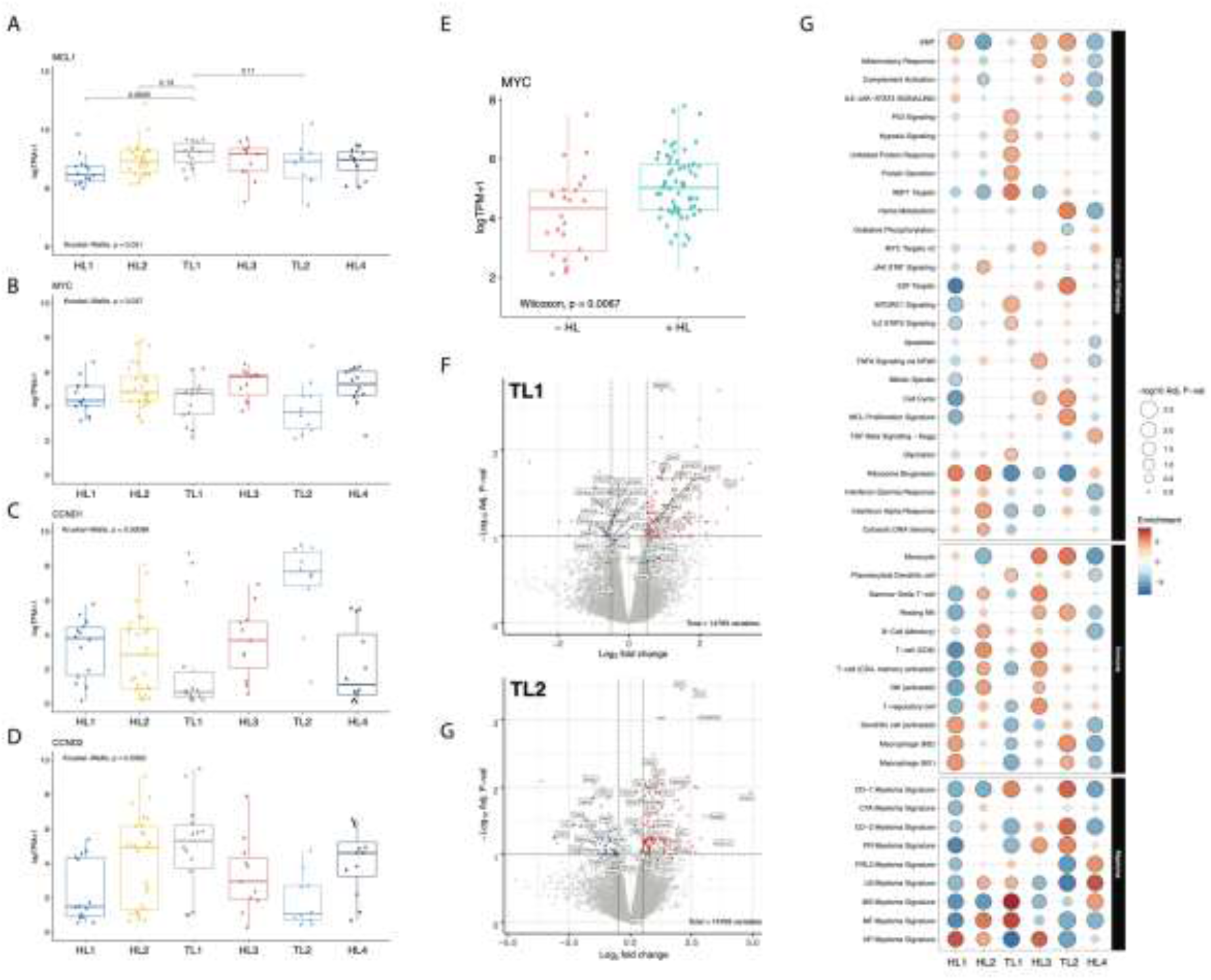
**A-D**) Comparison the expression levels of *MCL1, MYC, CCND1*, and *CCND2* genes among the six genetic subtypes.. **E)** Comparison the expression levels of *MYC* oncogene between the four hyperdiploid (HP) subgroups and the two non hyperdiploid ones. Expression is measured by the log value of transcript per million of each gene, and the comparison is done using the Kruskal-Wallis test. **F)** Differential expression (DE) analysis of *FMD* genetic subtype vs. the others. **G)** DE analysis of *CND* genetic subtype vs. the others. **H)** Gene set enrichment analysis of different molecular and oncogenic pathways (top), immune cell signatures (middle), and MM-specific signatures (bottom) among the six genetic subtypes.

### The genetic subgroups are enriched with specific MM expression signatures

To date, ten distinct RNA expression signatures have been defined and validated as prognostic in newly diagnosed and relapsed MM patients^14,15^. Each expression signature was then associated to specific primary genetic lesions identified by fluorescent in situ hybridization (FISH), including hyperdiploidy and IgH translocations that activate c-MAF and *MAFB, CCND1, CCND3*, or *MMSET*^14,15^. We asked whether these expression signatures were present in our SMM cohort and correlated with the six genetic subgroups. To address this, we performed a gene-set enrichment analysis of these expression signatures among the genetic subtypes (lower panel of **Figure 2G**). We observed that the hyperdiploid expression (HP) signature^14,15^, which is seen in hyperdiploid MM patients, is upregulated in the tumors of our hyperdiploid clusters (HL1-4). The Cyclin D(CD) expression signatures, including CD-1 that highly expresses *CCND1* and CD-2, which expresses the B cell markers *CD20, CD79A*, and *CCND1* were significantly upregulated in the *TL2* genetic subgroup. Moreover, the high-risk MMSET (MS) expression signature, which is enriched in patients with t(4;14) and upregulates *MMSET* and *FGFR3* genes, was upregulated in the *TL1* cluster. The MAF (MF) signature, which has been reported in patients with t(14;16) and t(14;20) that upregulate *MAF* and *MAFB* genes, respectively, was enriched in both the *TL1* and *HL2* subgroups, consistent with the presence of these genetic alterations in their tumors. The low bone (LB) disease signature, which has not been previously mapped to a specific MM genetic alteration, was upregulated in the *HL4, TL1*, and *HL2* subgroups, suggesting it could be linked to 1q gain, which occurs frequently in these three subgroups. Interestingly, the PR signature, which is found in proliferative tumor cells, was enriched in the *HL3* and *TL2* subgroups. Furthermore, the NFkB signature was upregulated only in *HL2*, which could be explained by the high frequency of 16q deletions and *CYLD* mutations in this subgroup. Finally, the PRL3 signature, which overexpresses the protein tyrosine phosphatase *PTP4A3* and 27 additional genes, was upregulated only in *HL4*. This suggests it could also be linked to the presence of 1q gain, which is found in all the tumors of the *HNF* subgroup.

We further examined whether our genetic subtypes were enriched in specific mutational signatures for 72 samples with matched normal whole exome sequencing (**Methods)**. We found that the APOBEC mutational signature activity (SBS 2,13 COSMIC v3.0) differed between the genetic subtypes (*P* = 0.027, Kruskal-Wallis) while AID mutational signatures did not (*P* = 0.17) (Supp Fig 1E-G). Specifically, we found APOBEC activity enriched in the *HL2* & *TL1* clusters vs. the rest of tumors (P = 0.006) (Supp Fig 1H).

### Genetic subgroups have distinct transcriptional profiles

We performed GSEA on the available transcriptomic resulting data to explore which genes and biological pathways were differentially expressed among the genetic subgroups we identified. Pathways that were significantly enriched within the six genetic subtypes are described and illustrated (**Figure 2G**). We found that protein secretion, unfolded protein response (UPR), glycolysis, hypoxia, and mTOR signatures were specifically enriched in the *TL1* subgroup, while E2F target genes, cell cycle, heme metabolism, complement, and proliferation signatures were enriched in *TL2* tumors. Genes induced by MYC were highly expressed in *HL3* and *HL4*, consistent with MYC upregulation in these two clusters. The NFkB, cytosolic DNA sensing, and JAK-STAT signatures were enriched in the tumors of *HL2*. The interferon alpha and gamma response signatures were high in *HL2* but low in *TL1*. Interestingly, oxidative phosphorylation, WNT-beta-catenin, and TGF-beta signaling were enriched only in tumors of *HL4*, and the TNFa and inflammatory signatures were uniquely enriched in *HL3*. Ribosome biosynthesis was low in *TL1, TL2*, and *HL3* but high in *HL1, HL2*, and *HL4* subgroups.

We also looked at signatures related to the tumor immune microenvironment. Signatures of regulatory T cells and NK cells were high in *HL2* and *HL3*, while the M2 macrophage signature was high in *TL2* and *HL4* tumors. The *HL3* and *TL2* tumors were enriched for the monocyte signature. In contrast, the signature of plasmacytoid dendritic cells, known for their immunosuppressive effect^16^, was only enriched in the *TL1* subtype.

### Genetic subtypes are differentially associated with risk of progression and evolving clinical biomarkers

To investigate the relationship between these genetic subtypes and clinical outcome, we analyzed a subset of patients (*n* = 87) who were followed for the natural course of their disease and did not receive any treatment in a clinical trial setting before progression to MM. Their baseline characteristics are reported in **Supp. Table 2**. The median follow-up time for these patients was 7.1 years and the median time to progression (TTP) was 4 years (95% CI, 3-6). In this cohort, 57 patients (66%) have progressed, while 30 (34%) remained asymptomatic as of the last follow-up (put date of last follow up in the methods section). The genetic subgroups had different outcomes, measured by TTP to active MM (log-rank *P* = 0.007) (**Supp Figure 7A**). Median TTP for patients in *HL2, TL1*, and *HL3* was 3.7, 2.6, and 2.2 years, respectively, while it was 4.3, 11, and 5.2 years for *HL1, TL2*, and *HL4*, respectively. The *HL2, TL1*, and *HL3* genetic subgroups had increased hazards of progression (HR > 4.5) to active myeloma (**Supp. Figure 7B**).

We then divided the genetic subtypes based on their TTP and hazards of progression into high-(*HL2, TL1, HL3*), intermediate-(*HL1, HL4*), and low-risk (*TL2*) subtypes. The high- and intermediate-risk subtypes had significantly shorter TTP and increased risk of progression compared to the low-risk subtype (2.6 and 5.2 vs. 11 years, respectively, *P* < 0.0001) (**Figure 3A**). We also stratified the patients according to the 20-2-20 model, which uses three cutoffs of M-protein > 2g/dL, FLC ratio > 20, and bone marrow plasmacytosis > 20% to define low, intermediate, and high-risk groups based on the presence of none, one, and two or all these variables, respectively^3^. The intermediate- and high-risk genetic subtypes and the clinically high-risk SMM group (according to the 20-2-20 model) were the only significant predictors of progression to active MM in our multivariate analysis (**Figure 3B**). Moreover, the prediction performance of the combined clinical and genetic models was higher than the clinical model alone (C-index: 0.76 vs 0.71, respectively) (**Supp. Table 3**). Interestingly, within the high-risk clinical group, patients in the high-risk genetic subgroups had increased progression risk (HR 3.7, 95% CI:1.1-12.8, *P* = 0.04). The two-year progression risk was 59% for the genetic high-risk compared to 41% for the genetic low-risk patients (**Supp. Fig 7C)**. We observed a similar findings in the intermediate-risk clinical group, where patients from the high-risk genetic groups had shorter TTP (3.4 vs. 9.4 years, *P* = 0.003, with a two-year progression risk of 33% vs. 0%, respectively). (**Supp. Fig 7D**). We found that high-risk genetic subtypes were significantly enriched with specific genetic alterations, such as, *KRAS, TP53*, t(4;14), t(14;16), 1q gain, 16q and 1p deletions among others (**Supp. Table 3**).

**Figure 3.**
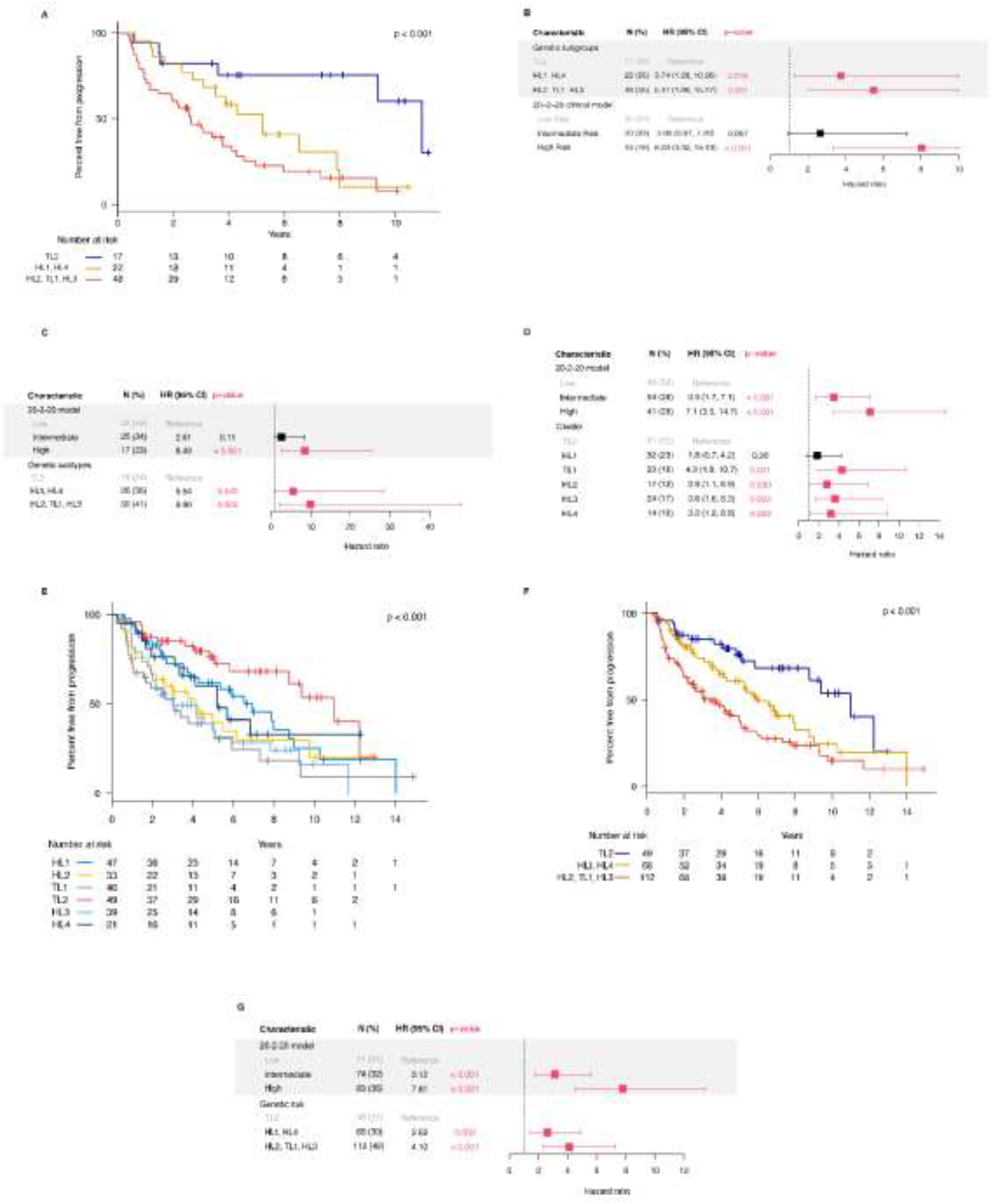
**A)** Kaplan-Meier curves for analysis of TTP in patients belonging to the three genetic risk groups. **B)** Multivariate cox regression analysis of the low, intermediate, and high-risk genetic subtypes and clinical risk stages according to the IMWG 20/2/20 model in the primary cohort. **C)** Multivariate cox regression analysis of the low, intermediate, and high-risk genetic subtypes and clinical risk stages according to the IMWG 20/2/20 model in the validation cohort. **D)** Multivariate cox regression analysis of the genetic subtypes and clinical risk stages according to the IMWG 20/2/20 model in the two validation cohorts of 74 and 67 patients. **E)** Kaplan-Meier curves for analysis of TTP in patients from the 6 genetic subtypes in the combined cohort of 229 patients. **F)** Kaplan-Meier curves for analysis of TTP in patients belonging to the three genetic risk groups of the combined cohort. **F)** Multivariate cox regression analysis of the low, intermediate, and high-risk genetic subtypes and clinical risk stages according to the IMWG 20/2/20 model in the combined cohorts.

We also identified patients with evolving M protein (eMP), which is defined as a ≥ 25% increase in M-protein within 12 months of diagnosis with a minimum absolute increase of 0.5 g/dL, and evolving hemoglobin (eHb), which is defined as a ≥ 0.5 g/dl decrease within 12 months of diagnosis^17^. These changing patterns were reported to confer a higher risk of progression to active MM in different SMM cohorts. We found that the odds of eMP and eHb were 9.4 and 5.3 times higher (*P* = 0.006 and 0.007, respectively) in patients with the high-risk genetic subtypes. These results indicate that high-risk genetic subgroups have distinct genetic and transcriptomic features as well as different clinical outcomes in terms of progression to active MM and evolution of its biomarkers over time.

### Validation of the molecular subtypes in external cohorts

To validate our findings on the clinical significance of the genetic subtypes, we developed a classifier based on the features of the clusters we identified in our primary cohort. We used an external cohort of 75 SMM patients to validate the classifier and investigate whether the genetic subtypes are predictive for progression^11^. The patients in this cohort were enriched in the low-risk clinical stage and had a median TTP of 5 years. Similar to the primary cohort, patients in the intermediate and high-risk genetic subtypes had increased risk of progression to active MM in multivariate analysis accounting for the clinical risk stage (HR: 4.5 and 9, *P* = 0.039 and 0.002, respectively) (**Figure 3C**). We found that adding the genetic risk groups improved the prediction of progression compared to the clinical model only (C-index: 0.76 vs 0.65, respectively) (**Supp. Table 4**). We also obtained another smaller cohort of 67 patients with targeted capture data, including common MM translocations, CNAs and SNVs, and added it to the previous cohort^12^. In those 142 patients, *HL2, TL1, HL3*, and *HL4* subtypes were independent predictors of progression to active myeloma (**Figure 3D**) and the high-risk genetic subtypes were associated with increased risk of progression in multivariate analysis (HR: 3.4, 95% CI :1.68-6.7). We then asked, given the small number of patients in the different cohorts, whether combining the three cohorts would provide more power and increase the significance of our genetic classification. The combined cohort contained 229 SMM patients with median follow-up and TTP of 6.9 and 5.2 years, respectively. Indeed, the genetic subtypes had a different TTP (**Figure 3F**), and the high-risk genetic subtypes had significantly shorter TTP compared to the low or the intermediate risk groups (**Figure 3F**). We also found that both the individual genetic subtypes and the genetic risk groups were independent predictors of progression in the combined cohort multivariate analysis, validating our initial findings (**Figure 3G**). We also observed that within the high-risk clinical stage, patients in the low-risk genetic subgroups had significantly lower progression risk (HR 0.26, 95% CI: 0.1-0.6, *P* = 0.001), while in the intermediate-risk clinical group, patients from the high-risk genetic groups were more likely to progress to symptomatic MM (HR 4.4, 95% CI: 1.7-11.6, *P* = 0.002) (**Supp. Fig 7F-H**).

## Discussion

This study modeled the genetic heterogeneity seen in SMM by identifying genetic subtypes that correspond to phenotypic attributes and clinical outcomes, providing a deeper understanding of SMM pathogenesis. We and others have previously cataloged individual driver genetic aberrations in SMM and MM cohorts^4,11,12^. However, the present study expands on this work and identifies SMM genetic subtypes defined by multiple recurrent DNA genetic aberrations, unlike previous classification efforts that were mainly based on gene expression data. Our findings suggest that these genetic subtypes could have distinct evolutionary histories depending on the initiating genetic events (translocations or CNAs), which may influence the subsequent acquisition of cooperating genetic aberrations.

The genetic subtypes had distinct clinical outcomes of disease progression into symptomatic MM, which could provide us with comprehensive molecular models for predicting progression and dynamic changes in clinical biomarkers over time. They also have specific dysregulated molecular and oncogenic pathways, which could facilitate the identification of specific targets and selection of therapies for each genetic subtype to empower precision medicine efforts, much like the efficacy of venetoclax specifically in patients with t(11;14).^18,19^

We identified six clusters based on the detected genetic alterations. We divided them into three high-risk (*HL2, TL1, HL3*), two intermediate-risk (*HL1, HL4*), and one low-risk (*TL2*) genetic groups based on progression risk to active MM. We found that DNA repair aberrations were exclusive to *HL2* and *TL1* subgroups, which were enriched in *TP53* mutations and deletions. Also, *MYC* expression was higher in the hyperdiploid subgroups than the non-hyperdiploid ones, consistent with previous reports of a higher frequency of *MYC* alterations in hyperdiploid MM patients^20^. The key Cyclin D genes, *CCND1* and *CCND2*, were highly expressed in *TL2* and *TL1*, respectively, while *MCL1* expression was not different among the genetic subtypes but was lowest in *HL1* tumors. Of note, *CCND1* and *CCND2* expressions were reported to distinguish between hyperdiploid groups, indeed in our four hyperdiploid clusters, we found the former to be enriched in tumors with 11q gain, while the latter is highly expressed in tumors without 11q gain. However, we couldn’t assess their prognostic impact due to the small number of samples with gene expression data in patients who were followed for their disease course.

The gene expression signatures of specific molecular and oncogenic processes also varied significantly between the genetic subgroups. For example, *TL1* tumors showed specific enrichment for protein secretion, ER stress, UPR, glycolysis and mTOR signaling. This molecular phenotype manifested clinically where patients with this genetic subtype had the highest increase in M-protein levels at six and twelve months from diagnosis. Such patients may benefit from drugs inducing cellular stress, such as proteasome inhibitors or novel molecules targeting the ER stress and UPR pathways^21^. Alternatively, *TL2* tumors were uniquely enriched with genes related to B-lymphocytes, cell cycle, heme metabolism, and complement activation signaling. Clinically, these patients had the longest TTP, lowest baseline M-protein level and the least increase over time. We also found that the *HL2* tumors were enriched for interferon alpha response, cytosolic DNA sensing, and JAK-STAT signatures. These results underscore the phenotypic difference among the genetic subtypes and provide a conceptual framework for further functional studies that aim to validate or therapeutically target the dysregulated pathways and tumor dependencies in them.

In our multicenter cohort, we found that the genetic subtypes also differed in the clinical outcome of progression to active MM. The three high-risk subgroups (*HL2, TL1*, and *HL3*) had a higher rate of progression and were associated with evolving hemoglobin and M-protein levels, showing that these subgroups are also predictive of the dynamic changes in MM clinical biomarkers over time. The high-risk genetic subtypes were independent risk factors of progression to overt MM after accounting for the clinical risk stage by the 20-2-20 model. Moreover, among those patients considered high- and intermediate-risk by this model, those with the high-risk genetic subtypes progressed faster to active myeloma than the rest in the same clinical risk group. Finally, to validate and test the significance of the genetic subtypes, we trained a classifier and tested it on two external SMM cohorts and found the genetic subtypes and risk groups to be predictive for progression in those external cohorts similar to the primary cohort. Furthermore, to increase the power of our analysis, we combined the three cohorts together and found the same effect with more significance levels compared to our initial findings. Of note, the genetic features enriched in the high-risk genetic group were also found to confer a higher-risk of progression as individual features, with exception of t(14;16) and t(14;20) (**Supp. Tables 5 and 6**). In fact, we and others haven’t found them to confer a high risk of progression on their own^4,10,11^. However, multiple studies have shown that t(14;16) is frequently associated with APOBEC signature and genomic instability ^4-10^. In our study it was found in 5% of patients and with similar rates in the validation cohorts, so larger studies with cohorts enriched for t(14;16) may be needed to confidently determine their prognostic significance in SMM. One of the limitations of our study is that we couldn’t assess the prognostic impact of the MM signatures in comparison to our DNA subgroups because of the small number of cases with this data. Moreover, we propose this genetic classification to be applied only in the SMM stage as we haven’t tested its prognostic significance in active or relapsed MM settings.

In conclusion, these findings move us closer to identifying the SMM patients who are truly at a high risk of disease progression through better predictive models that integrate the molecular makeup of the tumor cells and may also guide precision medicine efforts to match targeted therapies with the optimal patient groups in multiple myeloma asymptomatic stages.

## Methods

We studied a total of 214 patients with SMM at the time of diagnosis from multiple centers in the US and Europe. We performed whole-exome sequencing (WES) on 166 tumors, including 72 matched tumor-normal samples and 94 tumor-only samples. RNA sequencing was performed on 100 of the 166 tumors. We also performed targeted capture and sequencing with an in-house MM gene panel on 48 additional samples. Patients who presented with MM symptoms at diagnosis, including hypercalcemia, renal impairment, anemia, or bone lytic lesions (CRAB), or had any myeloma-defining event were excluded from the analysis^22^. All samples were obtained after written informed consent, according to the Declaration of Helsinki. Fisher’s exact test was used to test for association between categorical variables. ORs and 95% CIs were calculated for binary outcomes from contingency tables or logistic regression for continuous predictors. The Wilcoxon or Kruskal-Wallis rank-sum test was used to assess a location shift in the distribution of continuous variables between two or more than two groups, respectively. Descriptive statistics (proportions, medians, etc.) were reported with 95% exact binomial CIs or range. Time-to-event endpoints were estimated using the method of Kaplan and Meier. Time to progression (TTP) was measured from the date of diagnosis to the date of documented progression to MM. Differences in survival curves were assessed using a log-rank test. Median follow-up was calculated using the reverse Kaplan-Meier method. Cox modeling was performed to assess the impact of specific variables on clinical outcome measures. All *P* values were two-sided, and adjustment for multiple hypothesis testing was performed using the Benjamini and Hochberg method^23^; *P* and q value thresholds for significance were set at 0.05 and 0.1. Statistical analysis is described in detail in the Supplemental Methods.

### BMF Clustering Workflow

To identify patients with shared, co-occurring DNA features, we applied a variant of non-negative matrix consensus clustering algorithm adapted for binarized input and output features, Binary Matrix Factorization (BMF). Our input matrix for subtyping consisted of a combined binarized input matrix of 42 driver genes, CNVs, and 5 translocations. To select the number of clusters (K) for the consensus clustering, we randomly downsampled our input matrix and computed silhouette scores using Dice dissimilarity, residuals of factorization fit, variance explained, and K-L divergence on binary matrix factorizations over a range of K. We found a decrease in K-L divergence with our full dataset from K = 5 to K = 6, which suggested that 6 clusters were best suited to ensure a converged factorization for N = 214 (Supp. Fig 2A-E). Additionally, we found that variance stabilized when we performed down sampling analyses at N = 75-100, suggesting we were powered to perform binary matrix factorization for a cohort at this minimum size. We concluded that a minimum of 100 samples and 6 clusters were suited for this approach. We take the following steps for subtyping:

1. Run BMF for our primary cohort (n=214) from K=2 to K=10
2. Run hierarchical clustering of the consensus matrix with Euclidean distance and Ward linkage
3. Select K=6 clusters from downsampling results

We assessed binary feature importance by performing a Fisher’s exact test to count feature representation within each cluster and outside of this cluster, testing for an equal proportion. The false discovery rate (FDR) was calculated using the Benjamini-Hochberg procedure.

### Subtype Classifier

We trained a random forest classifier on 36 overlapping translocations, SNVs, and CNVs between both our primary cohort and validation cohort found in at least 3 or more patients to predict molecular subtypes for each patient. We used sklearn’s Random Forest Classifier class and reported a mean 5-fold cross validation accuracy on our primary cohort of 86.7% (SD +/-5%) after performing a randomized grid search to hypertune parameters. The classifier was then used on unseen data on 75 SMM samples.

### Bulk RNA-Sequencing

We processed a subset of 89 matched RNA samples out of the 214 patients using the GTEx V8 pipeline^26^, aligned to Hg19 and using the Gencode v19 gene annotation.

### RNA Differential Expression and Pathway Analysis

We performed one vs. rest gene differential expression for each identified DNA-based subtype in matching RNA samples. The limma-voom pipeline^27^ was used with FDR computed using the Benjamini-Hochberg procedure. We performed ranked gene-set enrichment analysis (GSEA) using the fGSEA R package, with a rank of signed-log fold-change from limma-voom. We computed pathway enrichments for the HALLMARK and KEGG gene sets from MsigDB^28,29^.

### Mutational Signatures

We use default settings of SignatureAnalyzer to extract *de novo* mutational signatures from a 96 base-pair context for 72 samples with WES. Extracted signatures were mapped to Cosmic 3.0 using cosine similarity.

## Code Availability

All code is provided for reproducibility: https://github.com/getzlab/SMM_clustering_2020

## Data Availability

The DNA and RNA sequencing data and analyses presented in the current publication have been deposited in and are available from the dbGaP database under dbGaP accession phs001323.v2 p1.

## Author Contributions

M.B, S.A, F.A, I.M.G, and G.G conceived the study. M.B, S.A, F.A, R.S.P, R.R, E.K, J.P, C.J.N, S.C, and L.B collected the data. M.B, K.S, T.H.M, M.R, C.J.N, A.D, and C.J.B performed the experiments and prepared DNA libraries. M.B, S.A, F.A, R.S.P, R.R, B.Z, A.J.D, L.T, C.S, I.M.G, and G.G analyzed the data. M.B, S.A, F.A, R.S.P, R.R, G.G, and I.M.G wrote the manuscript. All authors contributed to the scientific discussion, reviewed and edited the manuscript, and agreed on its content.

## Acknowledgments and Funding

We would like to acknowledge Anna Justis, PhD for scientific writing support and Jean-Baptiste Alberge for critical review of the manuscript.

This study was supported in part by the National Institutes of Health (NIH; R01CA205954), the Multiple Myeloma Research Foundation (MMRF) Prevention Project-Perelman Family Foundation Early Disease Translational Research Program, a Leukemia and Lymphoma Society (LLS) SCOR grant, the Dr. Miriam and Sheldon G. Adelson Medical Research Foundation, and a Cancer Research UK (CRUK) Early Detection and Diagnosis Program award. Selina Chavda receives funding from the Medical Research Council, UK. Kwee Yong receives funding from the UCLH NIHR Biomedical Research Centre.

This research was also supported by a Stand Up To Cancer Dream Team Research Grant (Grant Number: SU2C-AACR-DT-28-18). Stand Up To Cancer is a program of the Entertainment Industry Foundation. Research grants are administered by the American Association for Cancer Research, the scientific partner of Stand Up To Cancer. Opinions, interpretations, conclusions, and recommendations are those of the author(s) and are not necessarily endorsed by Stand Up To Cancer, the Entertainment Industry Foundation, or the American Association for Cancer Research.

We thank all the patients and their families for their contributions to this study.

