## Supplementary material for "Genetic Subtypes of Smoldering Multiple Myeloma are associated with Distinct Pathogenic Phenotypes and Clinical Outcomes": Data Supplement

### Data Supplement of:

**Supplemental Methods and Materials:**

**Patients.** We used next-generation sequencing technologies to study 214 patients with SMM at time of diagnosis with a total of 223 samples, including 5 serial samples. We performed whole exome sequencing (WES) of 72 matched tumor-normal samples (mean target coverage 109X), WES on 94 tumor-only samples (with mean coverage 174X), and targeted deep sequencing on 48 samples (mean target coverage 774X). Samples were collected at Dana-Farber Cancer Institute, University College London, Mayo Clinic, and the University of Athens in Greece, in addition to multiple centers in the US and Europe participating in clinical trial NCT02316106. For 4 cases, we obtained serial samples at the time of SMM diagnosis and time of progression to MM, and 1 case, who has not progressed to date, we sampled twice at the SMM stage. FISH data was used to determine the presence of IgH translocations. After approval of the study protocols by the institutional review boards and ethics committees of the participating institutions, samples were obtained after written informed consent according to the Declaration of Helsinki.

**Whole exome sequencing**. Tumor DNA was extracted from CD138+ cells from patients' bone marrow. For germline control (normal), DNA was obtained from buccal mucosa (saliva), or peripheral blood mononuclear cells. Genomic DNA was extracted using QIAamp DNA mini kit (QIAGEN) according to the manufacturer's protocols, and double-stranded DNA concentration was quantified using PicoGreen dsDNA Assay kit (Life Technologies). Libraries were prepared by Agilent SureSelect XT2 Target Enrichment kit. To capture the coding regions, we used the SureSelect XT2 V5+UTR capture probes (Agilent). All sequencing was performed on the Illumina HiSeq 4000 platform at the Broad Institute. For tumor only samples (n= 94), libraries were prepared and hybridized using Agilent SureSelect XT2 V5 capture probes (Agilent) and sequenced on Illumina HiSeq 2500 platform.

**Targeted deep sequencing.** Genomic DNA was extracted using QIAamp DNA micro kit (QIAGEN) according to the manufacturer's protocols. The libraries for targeted sequencing were prepared using SureSelect XT Reagent Kits (Agilent), and an in-house bait set targeting 117 genes, including pan-cancer driver genes and frequently mutated genes in MM. The libraries were quantified using Agilent Tapestation, then pooled and loaded onto the Illumina HiSeq 4000 sequencer.

**Computational analysis.** The output from Illumina software was processed by the Picard data processing pipeline to yield BAM files containing well-calibrated, aligned reads. We have utilized the Broad Institute and the Getz Lab CGA WES Characterization pipeline [https://github.com/broadinstitute/CGA_Production_Analysis_Pipeline] developed at the Broad Institute to call, filter, and annotate somatic mutations and copy number variation. The pipeline employs the following tools: MuTect[1], ContEst[2], Strelka[3], Orientation Bias Filter[4], DeTiN [5], AllelicCapSeg[6], MAFPoNFilter[7], RealignmentFilter, ABSOLUTE[8], GATK[9], PicardTools[10], Variant Effect Predictor [11], Oncotator [12]. Recurrent sCNAs were identified using the GISTIC2.0 algorithm [13]. We applied ABSOLUTE to estimate sample purity, ploidy, and absolute somatic copy numbers. These were used to infer the CCFs of point mutations from the WES data, following the framework previously described [8]. Mutations were thereafter classified as clonal based on the posterior probability that the CCF exceeded 0.90 and subclonal otherwise. Mutational signatures were identified using Signature Analyzer and mapped to COSMIC v3 reference signatures (Kim et al., 2016).

**Germline filtering of tumor-only cohort**

For each SNP or indel that passed all standard filters, its CCF, purity, ploidy, and local copy number were used to determine the log ratio of the probability that its allele fraction is consistent with the allele fraction modeled for a hypothetical germline event and the probability it is consistent with a modeled somatic event, as previously described (Chapuy et al., 2018). After applying the Germline Somatic Log odds filter, we used the ExAC database to further exclude potential germline events that have occurred in 5 or more participants of any ethnic background.

#### Artifact filtering

Both cohorts were subjected to standard artifact filtering through the Broad Institute's CGA pipeline, including a TCGA panel of normals (PoN) filter for common germline mutations and artifacts and filters for OxoG and FFPE damage.

##### *Paired tumor-normal cohort*

We applied ABSOLUTE to estimate sample purity, ploidy, and absolute somatic copy numbers. These were used to infer the CCFs of point mutations from the WES data. We excluded 13 samples from this group due to low tumor fraction (>20%) and inconclusive FISH results.

Bleed-through error (BTE) associated with sequencing was observed and cleaned using a custom panel of normals (PoN) run through the same sequencing pipeline, as described previously [7].

Two more artifacts were identified in this cohort, primarily characterized by C>A and C>T substitutions, respectively, henceforth referred to as A1 and A2. Artifact A1 was shown to represent reference bias, with a preponderance of C>A over G>T substitutions, related to oxidative damage occurring during DNA library preparation, as previously described [4]. To address this, we developed a tool that removes C>A SNPs with a low number of reads supporting the alternate allele from a sample, until the p-value of a binomial test assuming a probability of 0.5 exceeds 0.1. Of unidentified origin, artifact A2 was characterized by a preponderance of C>T SNPs in the GCC trinucleotide context over COSMIC signature 5 and was addressed by removing C>T SNPs in the GCC context with low alternate allele counts until they matched the number of C>T SNPs in the CCG context, assuming reference COSMIC signature 5 as a null. Of note, these artifacts did not affect any of the SNPs reported in genes that are frequently mutated in MM.

##### *Tumor-only cohort*

In this cohort, we observed an artifact of unidentified origin that was primarily characterized by T>G substitutions (supplemental figure, panel C) and hotspot mutations in genes never before reported in multiple myeloma, henceforth referred to as A3. We addressed it by removing all hotspot mutations with less than 2 occurrences in COSMIC and do not affect genes reported to be recurrently mutated in multiple myeloma. Of note, this artifact did not affect any of the SNPs reported in frequently mutated genes in MM.

**NMF Clustering Workflow**

We identified patient subgroups using binarized DNA features and performed consensus binary matrix factorization. To select K for the consensus clustering, we randomly down sampled our input matrix and computed silhouette scores using Dice dissimilarity, residuals of factorization fit, variance explained, and K-L divergence on binary matrix factorizations over a range of K. We found a decrease in K-L divergence with our full dataset from K=5 to K=6, which suggested that 6 clusters were best suited to ensure a converged factorization for N=214. Additionally, we found that variance explained stabilized when we performed down sampling analyses at N=75-100, suggesting we were powered to perform binary matrix factorization for a cohort at this minimum size. We concluded that a minimum of 100 samples and 6 clusters were suited for this approach. We performed consensus clustering using a binary matrix factorization with K of 2 through 10, selecting the final 6 clusters based on hierarchical clustering of the consensus matrix with Euclidean distance and Ward linkage. We assessed binary feature importance by performing a Fisher's exact test to count feature representation within each cluster and outside of this cluster, testing for an equal proportion. FDR was performed using the Benjamini-Hochberg procedure.

**Bulk RNA-Sequencing**

Out of the 214 unique patient tumor DNA samples, there were 89 matching RNA samples. These samples were isolated using Qiagen RNA kit. Libraries were prepared using Illumina Total mRNA kit and submitted for sequencing on Hiseq 2500 machines. We computationally processed these RNA samples using the GTEx V8 pipeline and aligned them to Hg19 Gencode v19.

**RNA Differential Expression + Pathway Analysis**

We performed 1 vs. rest gene differential expression for each identified DNA-based subtype. The limma-voom pipeline was used with FDR performed using the Benjamini-Hochberg procedure.

Using genes at an FDR < 0.1, we performed ranked gene-set enrichment analysis (GSEA) using the fGSEA R package, using a rank of -log10(FDR) * signed-log Fold-Change. We computed pathway enrichments for the HALLMARK and KEGG gene sets from MsigDB [15].

**Statistical analysis:** Binary outcomes were reported as proportions with 95% exact binomial confidence intervals. Continuous measures were summarized as median and range, and categorical variables were summarized as proportions. Binary outcomes and other categorical variables were tested for association with continuous and other categorical variables using Wilcoxon rank-sum (or Kruskal-Wallis for three or more groups) or Fisher's exact tests, respectively. Time-to-event endpoints are estimated using the method of Kaplan and Meier, with 95% confidence intervals calculated using Greenwood's method of variance estimation. Differences in survival curves were assessed using log-rank tests. Median follow-up was calculated using the reverse Kaplan-Meier method. Unadjusted and adjusted Cox modeling was performed to assess the impact of the presence of a MM driver on clinical outcome measures, alone and in the presence of clinical features known to impact outcome.

Time to progression (TTP) was measured from date of diagnosis to date of documented progression to MM. All P values were two-sided, and adjustment for multiple hypothesis testing was performed using the method of Benjamini and Hochberg; P and q value thresholds for significance were set at 0.05 and 0.1, respectively. Statistical analyses were performed using R version 3.6.0 (2019-04-26).

#### Tables and Figures with legends:

**Supplemental Table 1:** Baseline demographics and clinical characteristics of the 214 SMM patients.

| | Total<br>n = 214 (%) |
|---|---|
| **Age** | |
| Median (range) | 62 (34 - 85) |
| **Sex** | |
| Female | 115 (54) |
| Male | 99 (46) |
| **Race** | |
| Black or African American | 10 (5) |
| White | 204 (95) |
| **BM % involvement** | |
| Median (range) | 0.250 (0.028 - 0.800) |
| **M-spike** | |
| Median (range) | 1.80 (0.00 - 5.17) |
| **LDH** | |
| Median (range) | 174.5 (92.0 - 562.0) |
| *Missing* | *24 (11)* |
| **β2M** | |
| Median (range) | 2.40 (0.80 - 9.10) |
| **Mayo, 2008** | |
| Low | 48 (22) |
| Intermediate | 136 (64) |
| High | 30 (14) |
| **20-2-20 Risk score** | |
| Low | 53(25) |
| Intermediate | 63(29) |
| High | 98(46) |

**Supplemental Table 2:** Baseline demographics and clinical characteristics of the 87 untreated SMM patients who were followed for disease progression and their stratification according to the 20/2/20 clinical risk model.

| | | 20-2-20 clinical model | | | |
|---|---|---|---|---|---|
| | **Total** n = 87 (%) | **Low** n = 23 (27) | **Intermediate** n = 22 (26) | **High** n = 42 (55) | ***p-value*** |
| **BM % involvement** | | | | | |
| Median (range) | 0.30 (0.10 - 0.80) | 0.15 (0.10 - 0.20) | 0.20 (0.10 - 0.55) | 0.40 (0.20 - 0.80) | *< 0.001*† |
| **M-spike** | | | | | |
| Median (range) | 1.69 (0.00 - 5.17) | 1.18 (0.00 - 1.87) | 1.34 (0.00 - 2.30) | 2.16 (0.00 - 5.17) | *< 0.001*† |
| **FLCr, Involved/uninvolved ratio** | | | | | |
| Median (range) | 19.3 (1.1 - 325.3) | 2.9 (1.1 - 15.7) | 11.1 (1.6 - 255.9) | 48.6 (2.3 - 325.3) | *< 0.001*† |
| **β2M** | | | | | |
| Median (range) | 2.50 (1.40 - 6.40) | 2.50 (1.50 - 6.40) | 2.35 (1.50 - 3.80) | 2.50 (1.40 - 5.90) | *0.95*† |
| **Hemoglobin** | | | | | |
| Median (range) | 12.6 (8.3 - 16.2) | 12.7 (8.3 - 15.4) | 13.2 (11.0 - 16.2) | 12.4 (10.3 - 15.2) | *0.14*† |
| **LDH** | | | | | |
| Median (range) | 156 (92 - 422) | 147 (92 - 422) | 172 (120 - 262) | 160 (104 - 364) | *0.21*† |
| *Missing* | *15 (18)* | *6 (26)* | *7 (32)* | *2 (5)* | |
| **Calcium** | | | | | |
| Median (range) | 9.60 (8.70 - 10.50) | 9.60 (8.70 - 10.50) | 9.45 (8.80 - 10.00) | 9.60 (9.00 - 10.10) | *0.82*† |
| **Creatinine** | | | | | |
| Median (range) | 0.90 (0.60 - 1.60) | 0.90 (0.69 - 1.40) | 0.99 (0.60 - 1.60) | 0.90 (0.60 - 1.40) | *0.12*† |
| *Missing* | *1 (1)* | - | *1 (5)* | - | |
| **Total protein** | | | | | |
| Median (range) | 8.1 (5.5 - 11.0) | 7.7 (6.8 - 8.4) | 7.8 (5.5 - 8.8) | 8.6 (5.5 - 11.0) | *< 0.001*† |
| **Albumin** | | | | | |
| Median (range) | 4.1 (2.9 - 5.0) | 4.2 (3.5 - 5.0) | 4.2 (3.5 - 4.8) | 4.0 (2.9 - 4.8) | *0.10*† |
| *Missing* | *1 (1)* | *1 (4)* | - | - | |
| **Globulin** | | | | | |
| Median (range) | 3.8 (0.3 - 7.8) | 3.7 (2.2 - 4.6) | 3.5 (2.0 - 4.6) | 4.2 (0.3 - 7.8) | *0.003*† |
| *Missing* | *1 (1)* | *1 (4)* | - | - | |
| **Albumin/Globulin ratio** | | | | | |
| Median (range) | 1.1 (0.4 - 13.0) | 1.1 (0.8 - 2.3) | 1.2 (0.9 - 1.8) | 0.9 (0.4 - 13.0) | *0.005*† |
| *Missing* | *1 (1)* | *1 (4)* | - | - | |

†Cuzick's trend test

**Supplemental Table 3:** List of genetic alterations that were significantly enriched in High-risk genetic subgroups.

| Genetic alteration | pval | pval_adj |
|---|---|---|
| **16q_del** | 3.92133701244461E-19 | 1.88224176597341E-17 |
| **6q_del** | 1.30661737533785E-16 | 3.13588170081083E-15 |
| **del_22q** | 3.46052641707861E-15 | 5.53684226732578E-14 |
| **del_20q** | 9.81983940020603E-14 | 1.17838072802472E-12 |
| **KRAS** | 5.74309437412292E-12 | 5.09021145715801E-11 |
| **1p_del** | 6.36276432144751E-12 | 5.09021145715801E-11 |
| **del_8p** | 1.076934966262E-11 | 7.38469691151089E-11 |
| **17p_del** | 2.66210661752379E-10 | 1.59726397051427E-09 |
| **t(4;14)** | 9.59521520668869E-10 | 5.11744811023397E-09 |
| **14q_del** | 7.16829126182379E-09 | 3.44077980567542E-08 |
| **del_4q** | 1.03923060318069E-08 | 4.534824450243E-08 |
| **ATM** | 6.28534662885084E-08 | 2.32074337065262E-07 |
| **t(14;16)** | 6.28534662885084E-08 | 2.32074337065262E-07 |
| **DIS3** | 7.72727909076587E-08 | 2.64935283111973E-07 |
| **BRAF** | 5.07514260624449E-07 | 1.52254278187335E-06 |
| **LTB** | 6.5996139602957E-07 | 1.6672708952326E-06 |
| **HIST1H1E** | 6.5996139602957E-07 | 1.6672708952326E-06 |
| **t(14;20)** | 6.5996139602957E-07 | 1.6672708952326E-06 |
| **NFKBIA** | 7.64717173177041E-07 | 1.8353212156249E-06 |
| **del_10p** | 3.75087303233623E-06 | 8.18372297964268E-06 |
| **TP53** | 6.63104069343861E-06 | 1.32620813868772E-05 |
| **ZNF292** | 1.19077388394712E-05 | 1.97093608377455E-05 |
| **MAFB** | 1.19077388394712E-05 | 1.97093608377455E-05 |
| **1q_gain** | 3.91221254141483E-05 | 6.25954006626372E-05 |
| **FAM46C** | 4.98897812477866E-05 | 7.72486935449599E-05 |
| **MAF** | 8.10600203567709E-05 | 0.000111168027917857 |

**Supplemental Table 4:** Performance of the Clinical Models only and after adding the Genetic Model (based on the genetic risk groups).

| Cohort | Model | Likelihood Ratio Test Statistic | χ2 *P* | Model C-statistic† (95% CI) | AIC/BIC |
|---|---|---|---|---|---|
| Primary (DFCI, UCL, Greece) | Clinical | 14.7 | < 0.001 | 0.71 (0.64 - 0.78) | 406/411 |
| | Clinical + Genetic | | | 0.76 (0.68 - 0.83) | 396/404 |
| Validation (UAMS) | Clinical | 12.1 | 0.002 | 0.65 (0.49 - 0.80) | 165/168 |
| | Clinical + Genetic | | | 0.76 (0.62 - 0.90) | 157/162 |
| Validation (UAMS, Mayo) | Clinical | 13.53 | 0.001 | 0.66 (0.59 - 0.74) | 567/571 |
| | Clinical + Genetic | | | 0.71 (0.64 - 0.78) | 560/569 |
| Combined (Primary, UAMS, Mayo) | Clinical | 29.84 | < 0.001 | 0.69 (0.64 - 0.73) | 1146/1151 |
| | Clinical + Genetic | | | 0.74 (0.69 - 0.78) | 1123/1135 |

Improvement in goodness of fit was assessed with a likelihood ratio test. The genetic model significantly improved the fit of the clinical-only models. A global assessment of each model was also assessed using a C-statistic for censored survival data.10 The statistic for each time-to-event model is reported with a 95% CI. Values range from 0.5 to 1, which indicates a poor to perfect model.

**Supplemental Table 5: Univariate Cox PFS regression of the high-risk genetic features in the primary cohort (n = 87).**

| | | Estimates | | p-value | |
|---|---|---|---|---|---|
| **Variable** | **No. of patients (%)** | **HR** | **95% CI** | **naive** | **corrected** |
| MYC alterations | 7 (9) | 7.58 | 3.02 - 19.03 | < 0.001 | < 0.001 |
| KRAS | 12 (14) | 4.13 | 2.10 - 8.12 | < 0.001 | < 0.001 |
| del(1p) | 7 (8) | 4.24 | 1.89 - 9.51 | < 0.001 | 0.003 |
| del(14q) | 10 (12) | 3.06 | 1.47 - 6.34 | 0.003 | 0.010 |
| del(8p) | 8 (9) | 3.43 | 1.53 - 7.70 | 0.003 | 0.010 |
| TP53 | 5 (6) | 3.89 | 1.50 - 10.10 | 0.005 | 0.017 |
| NRAS | 5 (6) | 3.23 | 1.27 - 8.21 | 0.014 | 0.035 |
| del(6q) | 11 (13) | 2.46 | 1.18 - 5.11 | 0.016 | 0.035 |
| del(22q) | 7 (8) | 2.97 | 1.24 - 7.09 | 0.014 | 0.035 |
| t(4;14) | 5 (6) | 2.69 | 1.05 - 6.89 | 0.039 | 0.072 |
| del(16q) | 18 (21) | 1.66 | 0.89 - 3.08 | 0.11 | 0.06 |
| del(4q) | 5 (6) | 1.69 | 0.61 - 4.69 | 0.32 | 0.41 |
| del(13q) | 42 (49) | 1.26 | 0.73 - 2.17 | 0.41 | 0.50 |
| gain(1q) | 21 (25) | 1.6 | 0.90 - 2.83 | 0.47 | 0.12 |

**Supplemental Table 6: Univariate Cox PFS regression of the high-risk genetic features in the combined cohorts (n = 229).**

| Variable | No. of patients (%) | Estimates | | p-value | |
|---|---|---|---|---|---|
| | | HR | 95% CI | naive | corrected |
| MYC_aberrations | 48 (21) | 1.9 (1.2-2.8) | | 0.0022 | 0.0055 |
| KRAS | 43 (19) | 2.7 (1.9-4) | | 3.50E-07 | <0.0001 |
| gain_1q | 60 (26) | 1.5 (1-2.1) | | 0.047 | 0.0588 |
| del_16q | 38 (17) | 1.4 (0.91-2.2) | | 0.12 | 0.1385 |
| del_1p | 24 (11) | 2 (1.3-3.3) | | 0.0027 | 0.0058 |
| del_8p | 20 (9) | 3.2 (1.9-5.4) | | 1.10E-05 | 0.0001 |
| del_6q | 29 (13) | 2.6 (1.7-4.1) | | 2.10E-05 | 0.0001 |
| t.4.14. | 16 (7) | 2.6 (1.5-4.6) | | 0.00082 | 0.0025 |
| TP53_aberrations | 14 (6) | 2.6 (1.4-4.8) | | 0.0035 | 0.0066 |
| DIS3 | 13 (6) | 2.3 (1.2-4.3) | | 0.0082 | 0.0123 |
| FAM46C | 9 (4) | 2.7 (1.3-5.7) | | 0.0067 | 0.0112 |
| del_22q | 22 (10) | 1.7 (1-2.9) | | 0.039 | 0.0532 |
| del_14q | 23 (10) | 2.6 (1.6-4.3) | | 0.00011 | 0.0004 |
| t.14.16. | 6 (3) | 0.98 (0.35-2.7) | | 0.97 | 0.9700 |
| ATM | 6 (3) | 2.1 (0.75-5.6) | | 0.16 | 0.1714 |

**Supplemental Figure 1** A) Age and sex distribution of primary cohort (n=214). B) Logistic principal components analysis (PCA) projection of binarized dataset (n=214). Green color denotes the presence of the variable of interest while red color indicates the rest of the dataset. C) Sample-sample consensus matrix of binary matrix factorization results (K=2-10) with translocations and copy number (left). D) LogisticPCA colored by subtypes. E-F) Mutational signature abundance per subtype.

Figure S1

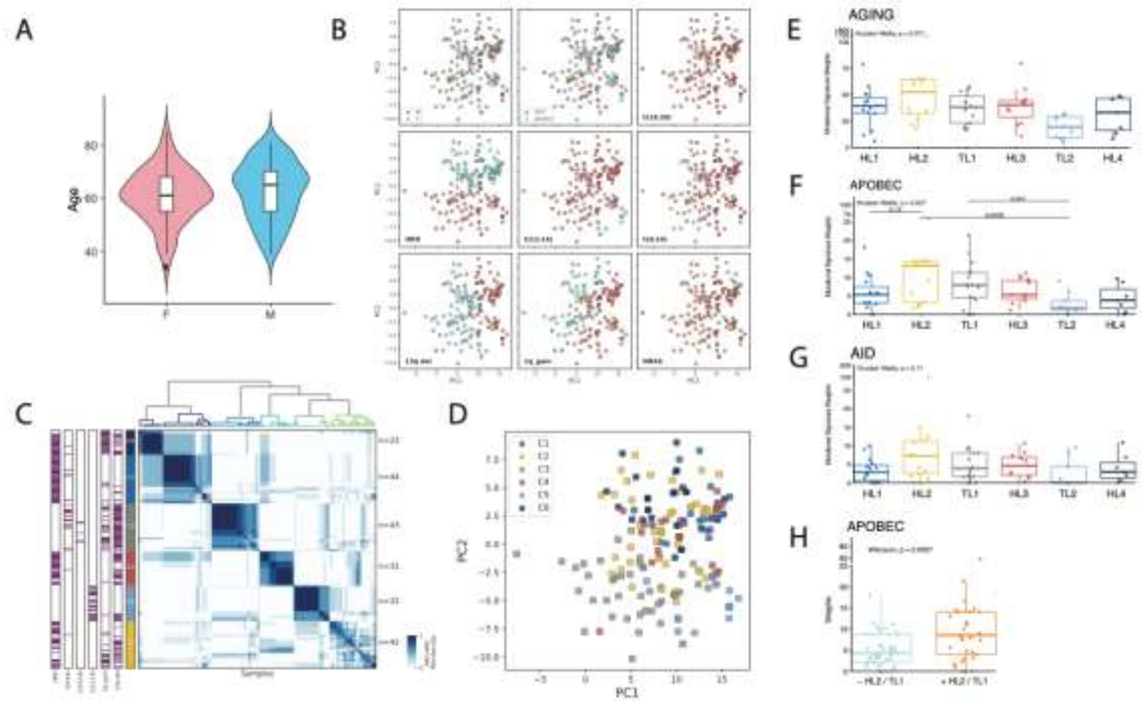

**Supplemental Figure 2:** Downsampling analysis of dataset for binary matrix factorization; A-D) random downsample runs (n=100) for a range of sample sizes (n=25-214) showing explained variance (A) K-L divergence (B) Residuals (C) and Silhouette Score (D); E) Heatmap of mean scores for each downsample run.

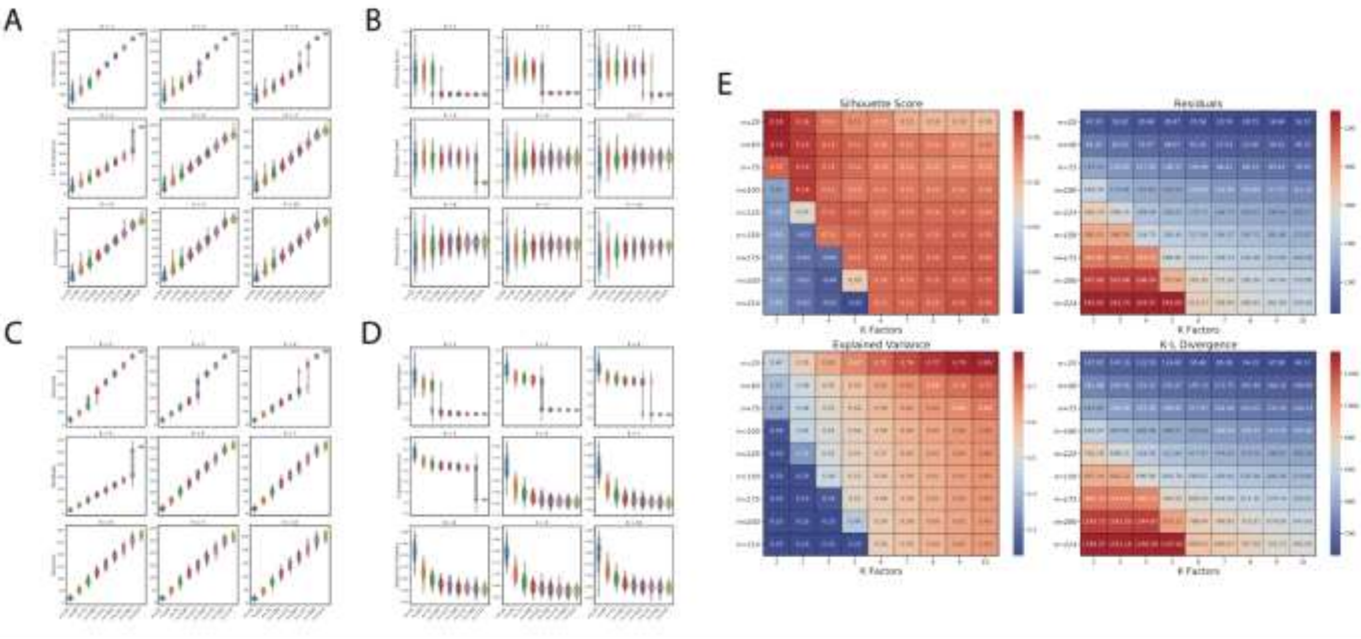

**Supplemental Figure 3:** RNA-SeQC quality control metrics for NGS RNA samples A) general metrics with filtered samples (n=13) for a cutoff of 0.8 median exon CV. B) PCA of filtered transcriptome colored by 3' Strand Bias C) Median Exon CV

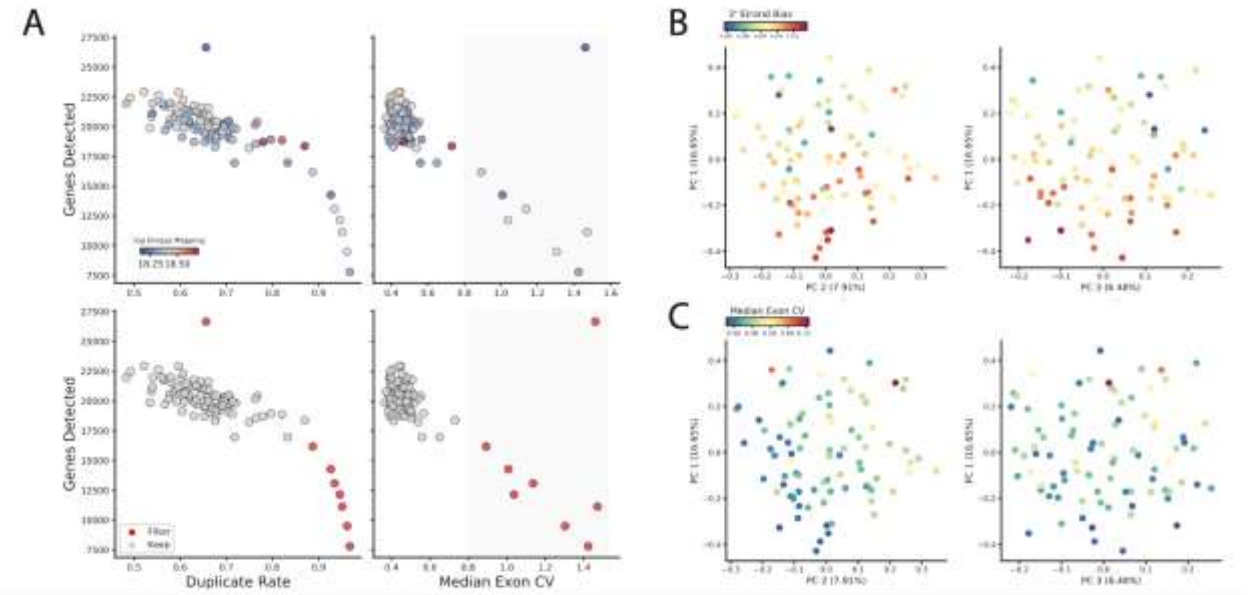

**Supplemental Figure 4:** Pathway enrichment enrichment based on log-fold change from Limma-Voom per subtype for **A)** KEGG B) Hallmark **C)** upregulated genes in myeloma signatures **D)** downregulated genes in myeloma signatures. **E)** Multiple myeloma gene expression signatures as mean log TPM among the six genetic subtypes for gene-sets combined from both Zhang et al [16] and Broyl et al [17]. **F)** Filtered pathway enrichment figure ranked using log-fold change * -log10(FDR). Significantly enriched pathways in certain genetic subtypes are circled in black.

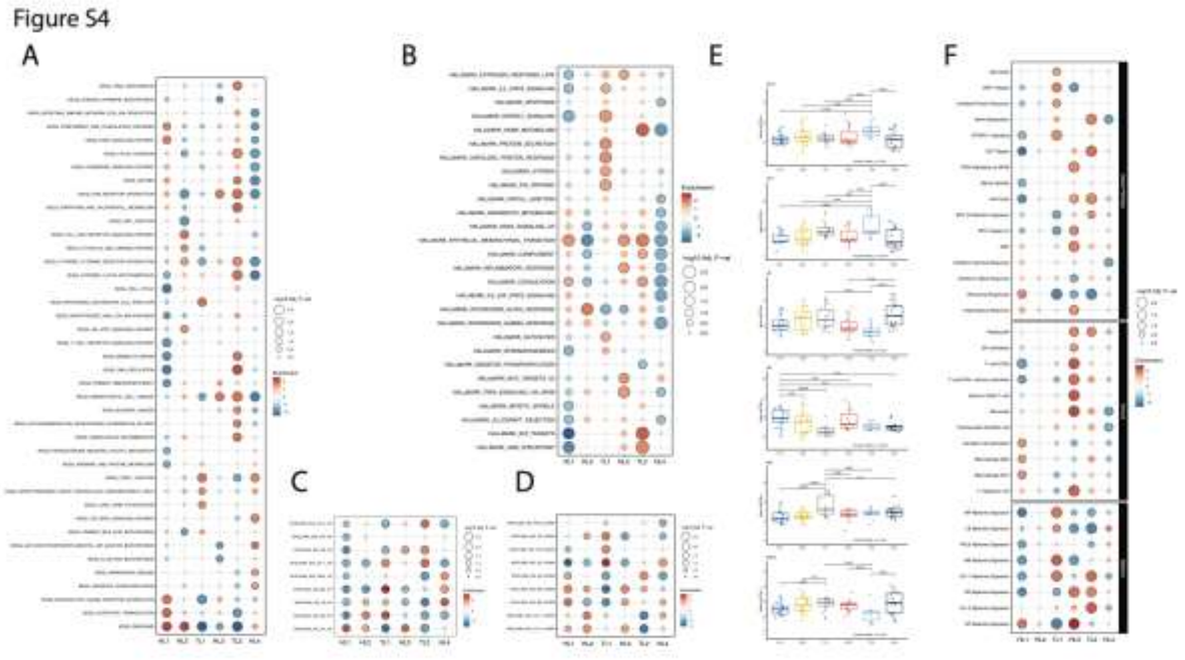

**Supplemental Figure 5: A)** Random forest K-fold cross validation accuracy over 10 repeats with mean accuracy and standard deviation reported. **B)** Grid search to optimize parameters did not improve over initial random forest parameters. **C)** The prevalence of each feature in the dataset plotted by feature importance in classifying genetic subtypes.

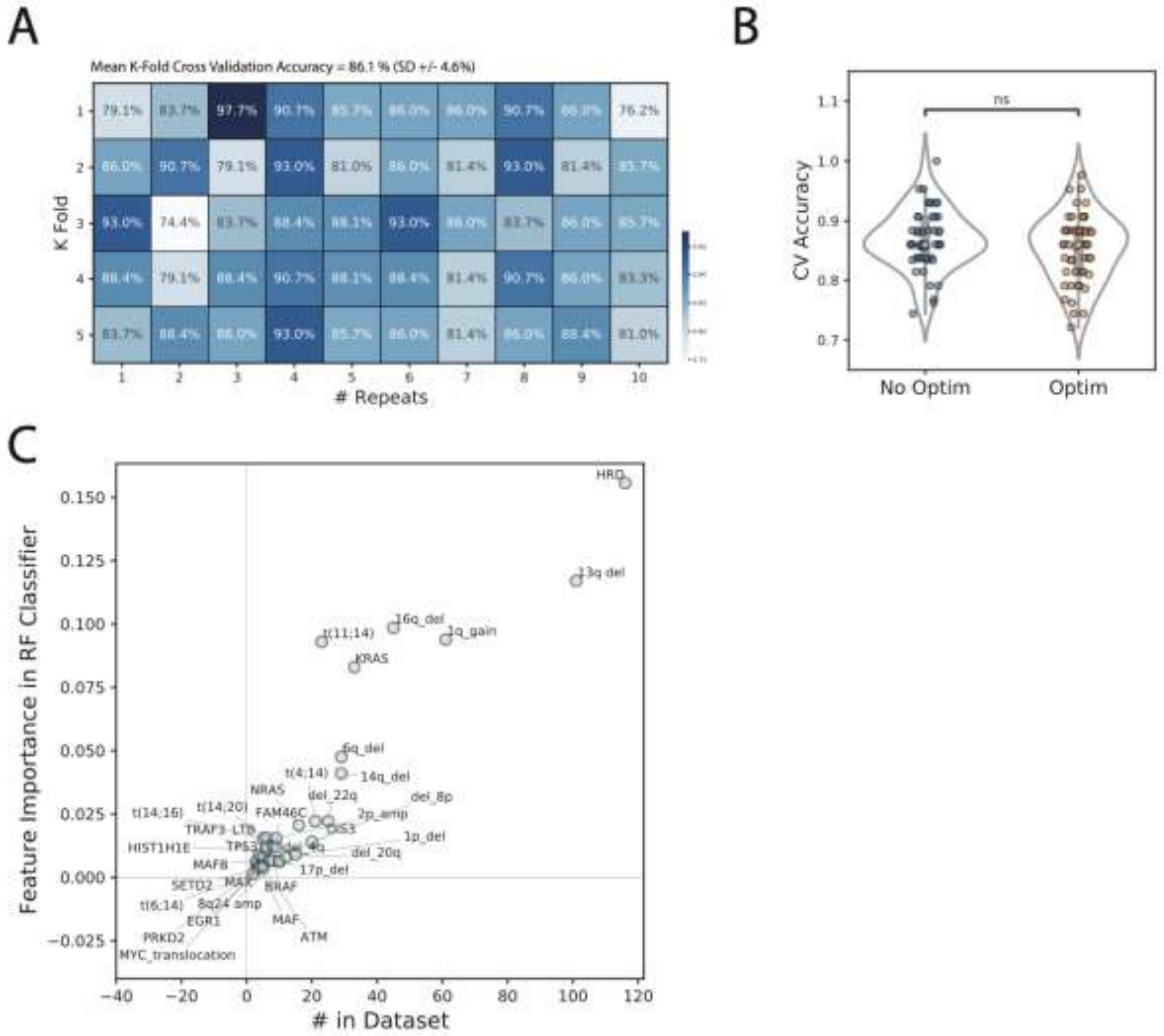

**Supplemental Figure 6:** Additional gene expression (log TPM + 1) comparisons. A) *CCND1* in TL2 tumors vs. the other tumors; B) *CCND2* in TL1 & HL2 tumors vs. the rest; C) *MCL1* expression in HL1 tumors vs. the rest. Within hyperdiploidy subtypes (HNT, HMC, HKR, HNF), further gene expression comparisons were done. D-F) *CCND1* expression in HNT, HMC, HKR, HNF subtypes with and without 11q gain, hyperdiploidy, and between these subtypes. G-I) *CCND2* expression in HNT, HMC, HKR, HNF subtypes with and without 11q gain, hyperdiploidy, and between these subtypes.

Figure S6

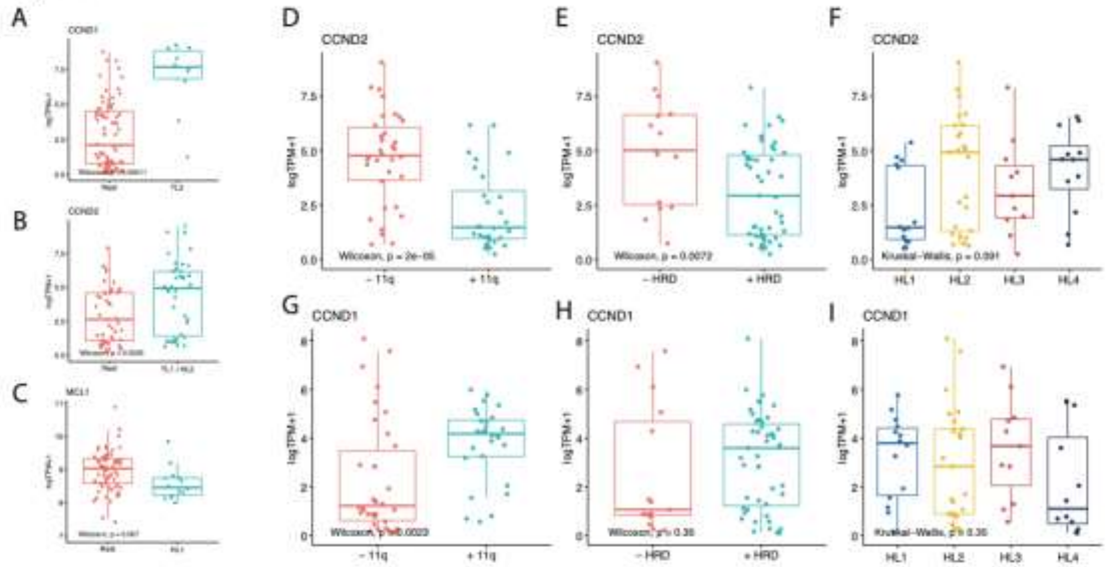

**Supplemental Figure 7:**

**A)** Kaplan-Meier curves for analysis of TTP according to the six genetic subtypes from the primary cohort. **B)** Cox proportional hazards analysis of the six genetic subtypes and in the primary cohort**. C)** Kaplan-Meier curves for analysis of TTP in patients from the primary cohort belonging to high vs intermediate and low-risk genetic subtypes in the clinically high-risk group by the 20-2-20 model. **D)** Kaplan-Meier curves for analysis of TTP in patients belonging to high vs intermediate and low-risk genetic subtypes in the clinically intermediate-risk group by the 20-2-20 model in the primary cohort. **E)** Kaplan-Meier curves for analysis of TTP in patients from the combined cohorts belonging to high vs intermediate and low-risk genetic subtypes in the clinically high-risk group by the 20-2-20 model. **F)** Kaplan-Meier curves for analysis of TTP in patients belonging to high vs intermediate and low-risk genetic subtypes in the clinically intermediate-risk group by the 20-2-20 model in the combined cohorts.

**A)**

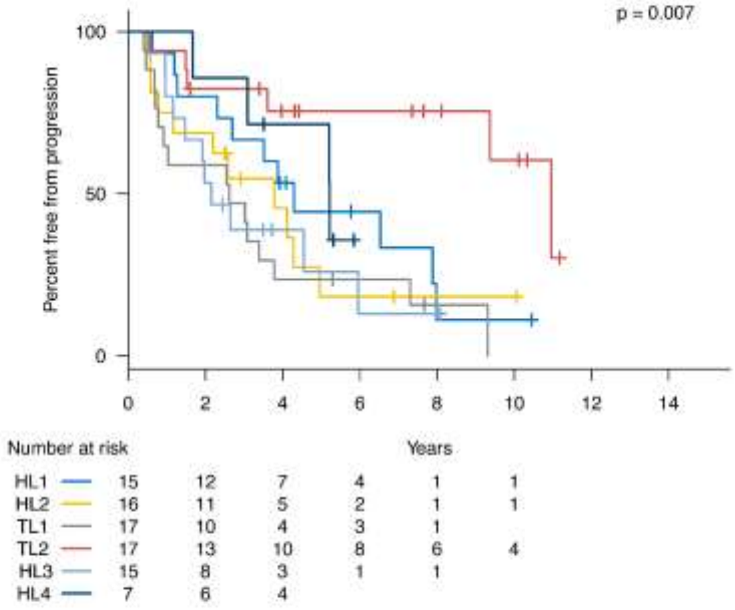
p = 0.007
Percent free from progression
100
50
0
0 2 4 6 8 10 12 14
Years
Number at risk
HL1 15 12 7 4 1 1
HL2 16 11 5 2 1 1
TL1 17 10 4 3 1
TL2 17 13 10 8 6 4
HL3 15 8 3 1 1
HL4 7 6 4

**B)**

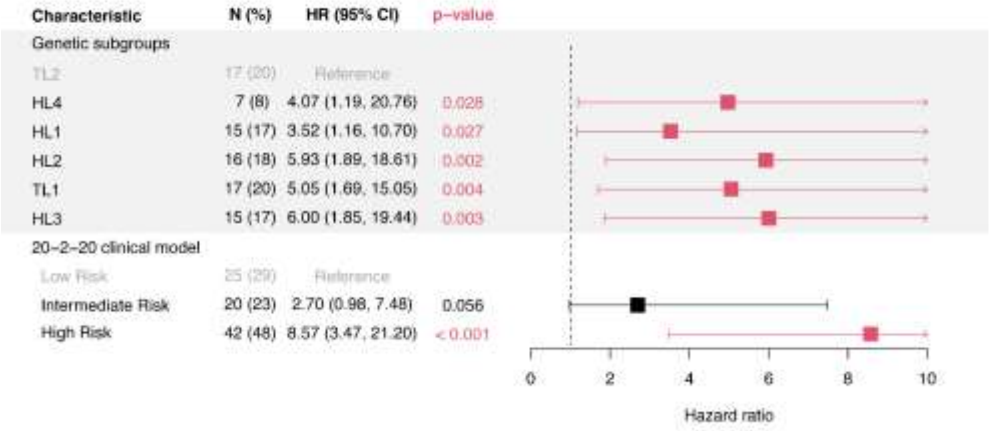
Characteristic
N (%)
HR (95% CI)
p−value
Genetic subgroups
TL2 17 (20) Reference
HL4 7 (8) 4.07 (1.19, 20.76) 0.028
HL1 15 (17) 3.52 (1.16, 10.70) 0.027
HL2 16 (18) 5.93 (1.89, 18.61) 0.002
TL1 17 (20) 5.05 (1.69, 15.05) 0.004
HL3 15 (17) 6.00 (1.85, 19.44) 0.003
20−2−20 clinical model
Low Risk 25 (29) Reference
Intermediate Risk 20 (23) 2.70 (0.98, 7.48) 0.056
High Risk 42 (48) 8.57 (3.47, 21.20) < 0.001
0 2 4 6 8 10
Hazard ratio

**C-E)**

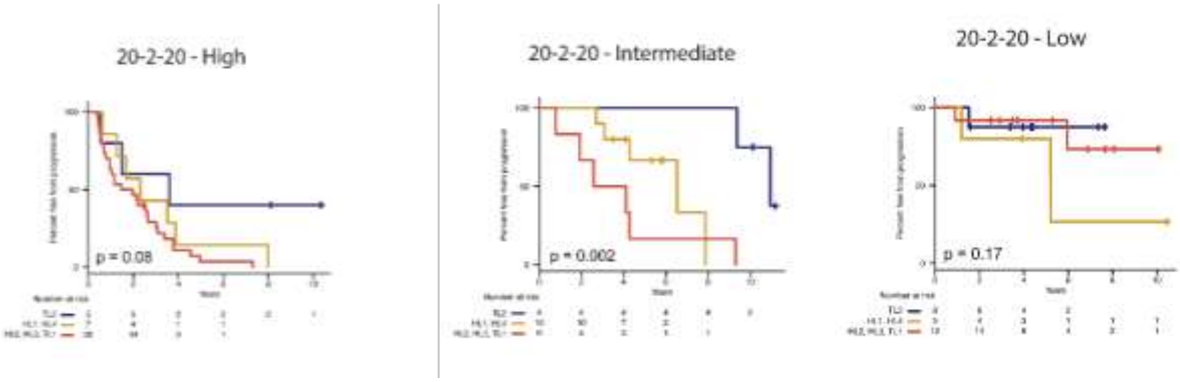
20-2-20 - High
p = 0.08
20-2-20 - Intermediate
p = 0.002
20-2-20 - Low
p = 0.17

**F-H)**

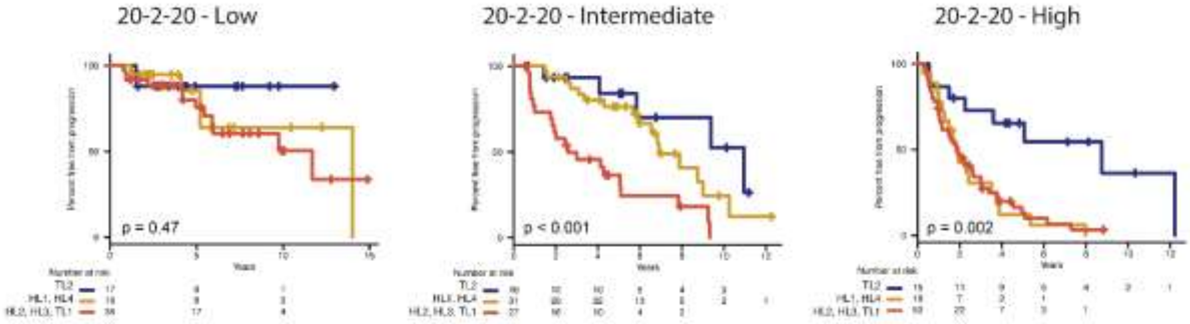
20-2-20 - Low
p = 0.47
20-2-20 - Intermediate
p < 0.001
20-2-20 - High
p = 0.002
TL2
HL1, HL4
HL2, HL3, TL1
